# Polymorphisms, Solvent Accessibility, and Evolutionary Conservation of Influenza A Virus PB1 Protein

**DOI:** 10.1101/2025.06.23.661066

**Authors:** Dejun Lian, Jie Lian, Qi Dong

## Abstract

Protein polymorphisms, reflecting amino acid and nucleotide sequence divergence, provide insights into protein evolution. Here, we analyze sequence variation and structural features of influenza A virus (IAV) PB1 protein, an RNA-dependent RNA polymerase critical for viral replication. Our findings demonstrate that residue solvent accessibility strongly predicts polymorphism likelihood, with exposed sites exhibiting higher variability. Despite extensive polymorphism, we observe pervasive purifying selection across PB1, maintaining functional and structural constraints. These results highlight how protein architecture shapes evolutionary dynamics in viral proteins.

## INTRODUCTION

Protein sequence divergence arises from mutations accumulating under evolutionary constraints. The neutral and nearly neutral theories posit that most molecular evolution stems from random fixation of neutral or mildly deleterious mutations, with purifying selection eliminating strongly detrimental variants (1,2,83). RNA viruses— including influenza, HCV, and SARS-CoV-2—exhibit exceptionally high mutation rates (10^−3^–10^−4^ mutations per nucleotide per replication), enabling rapid adaptation (3,4).

Influenza A virus (IAV), a major respiratory pathogen, evolves via mutation and genome reassortment, causing annual epidemics and pandemics. Its segmented genome encodes ten proteins, including PB1, the catalytic subunit of the RNA polymerase complex. PB1 lacks proofreading activity, resulting in high genetic diversity (8,9). While IAV evolution has been extensively studied (7-11), the mechanisms governing PB1 sequence variation remain incompletely understood.

Here, we investigate how PB1 structural features, particularly relative solvent accessibility (RSA), influence polymorphism patterns. RSA—quantifying residue burial within the protein core—predicts evolutionary rates across taxa (11). However, its generalizability to viral proteins warrants further exploration.

## Materials and Methods

### Sequences and Structures

A total of 107312 sequences of IAV isolates were retrieved from Genbank, Influenza 163 Research Database (IRD) and GISAID. The nucleotide sequences encoding the protein PB1 were used for comparative analysis. RNA Sequences were additionally filtered that only sequence length are larger than 1000 bases are accounted. IBV, ICV and IDV PB1 sequences analyzed in the study were also downloaded from GenBank. A listing of their accession numbers is available from the author upon request.

Sites that were variable within species were considered polymorphic, sites that were identical within species were treated as invariant sites. Only amino acids occurring more than once were counted, in order to avoid errors due to a single aberrant sequence.

The structures used in this study were determined through X-ray crystallography. We used PDB code 6r65 structure for analysis.

### Data analysis

Multiple protein and nucleotide sequences were aligned with BioEdit and edited by hand.

In our analysis, the SAS measure was used to estimate the proportion of each amino acid residue that is accessible to solvent. This was done by taking the ratio of SAS we calculated from the actual protein structure to that of the maximum exposed surface area in the fully extended conformation of the pentapeptide gly-gly-X-gly-gly, where X is the amino acid in question. We then normalized ASA values by the theoretical maximum ASA of each residue (14) to obtain RSA(relative solvent accessibility, solvent accessible surface area (SAS) of aa of PB1 are calculated using the DSSP program (http://www.cmbi.ru.nl/dssp.html) ( 16).

#### Statistical analysis

Data are expressed as means ± SD. Statistical analyses were performed using the Kruskal–Wallis and Mann–Whitney U methods. All statistical analyses were performed using SPSS version 13.0 (SPSS Inc., Chicago, IL) with additional analysis performed using Stata/MP14 (StataCorp LP). Values of p < 0.05 were considered significant.

### Logistic Regression, Confidence Intervals

We used the methods and model of Lian D (81) in understanding how a set of predictor variables affect a dichotomous outcome variable (polymorphic or invariant).

### The correlation of residue variability with structure

The correlation was tested using Lian D’s method (81). We performed several tests to find structural correlates of high evolutionary rate.

### The correlation of Entropy at position with structure

To measure the degree of sequence conservation, we used Lian D’s method.

### Ka/Ks

Ka/Ks values were obtained by Datamonkey Adaptive Evolution Server (http://www.datamonkey.org/). The multi-partition fixed effects likelihood (FEL) and FLAC method implemented in the Hyphy software package on the online server was then used to predict purifying selection. HCV sequences were clustered before used.

### McDonald and Kreitman test

McDonald and Kreitman test was performed using DnaSP 6 program. HCV sequences were clustered before used. We used Influenza B virus PB1 sequence as outgroup control.

### RNA structure prediction

MFED values were calculated by comparing minimum folding energies for WT and sequences shuffled in order by the algorithm NDR. Ensemble RNA structure predictions were made using the DAMBE program.

## RESULTS

### Maximum-Likelihood Estimates of Parameters for Logistic Regression of Polymorphism on Solvent Accessibility for Amino Acids Grouped by Protein

Table 1 summarizes the maximum-likelihood estimates of the parameters in the logistic regression model of polymorphism on solvent accessibility for amino acids. The second column lists the proportion of sites that vary within protein. The third and fourth columns give the maximum-likelihood estimates of α and βsas, respectively. The fifth column gives the results of likelihood ratio tests (LRTs) of whether the model with βsas 5 MLE(βsas) fits the data significantly better than a model with βsas= 0, where the test statistic is approximately distributed as χ^2^ with one degree of freedom.

**Table 1.**
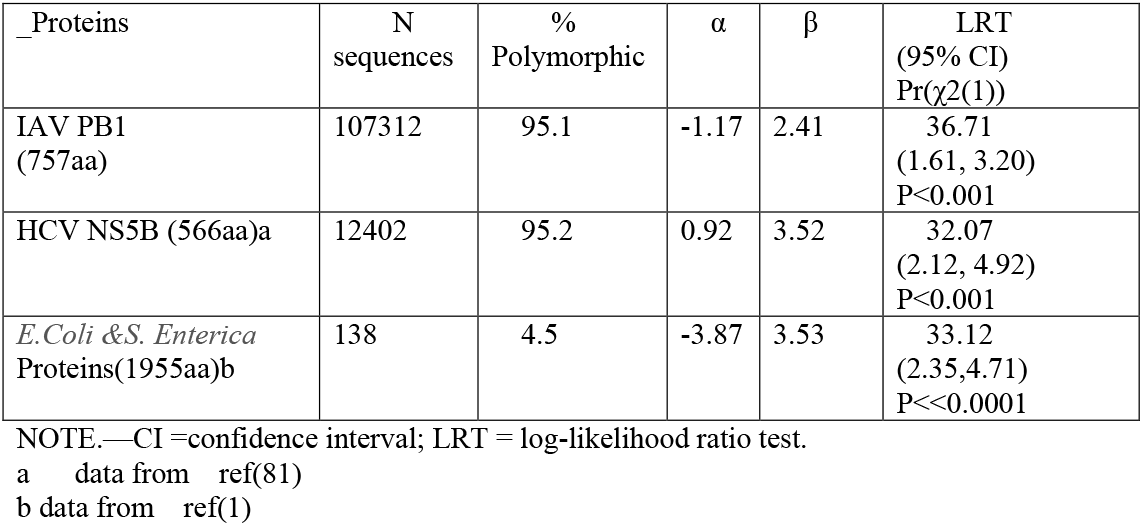
Logistic regression of aa polymorphism of PB1 on RSA.

### Residue variability

Residue variability (RV), a measure that roughly corresponds to states at residue sites. For IAV PB1 which gave a mean value of 3.4+-0.07, compares with HCV NS5B, 5.36+_0.14, and increase with RSA.

IAV PB1 is more conservative than HCV. >=4.4% sequences are monomorphic sites, a great many of the substitutions are conservative substitutions (the same physicochemical class)

**Fig.1.**
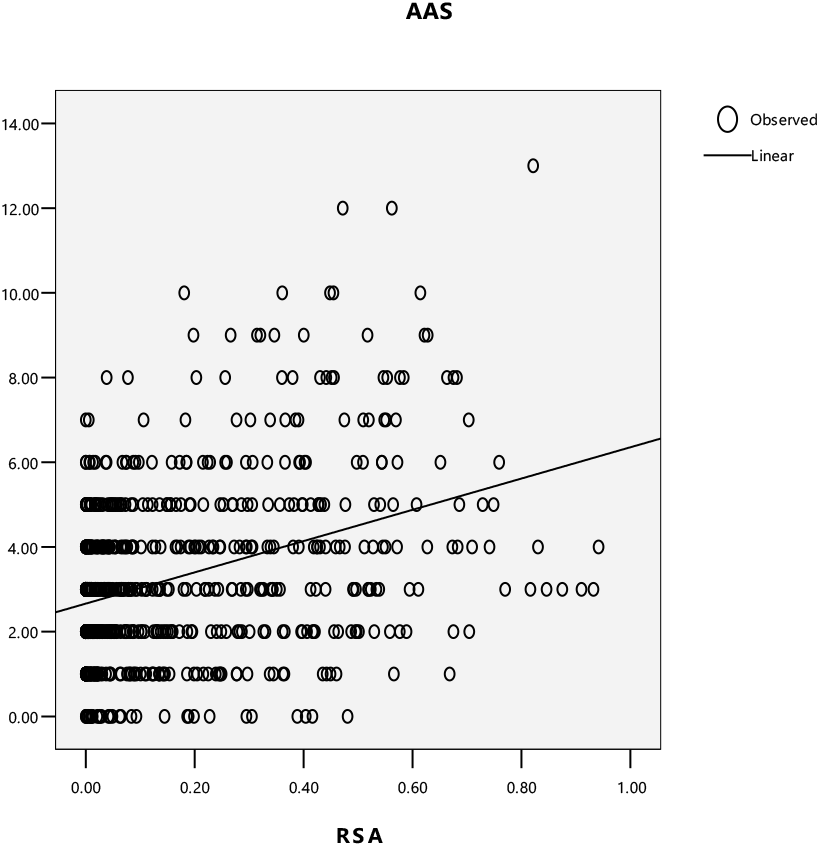
Liner regression of aa polymorphism of PB1 on RSA.

**Fig.2.**
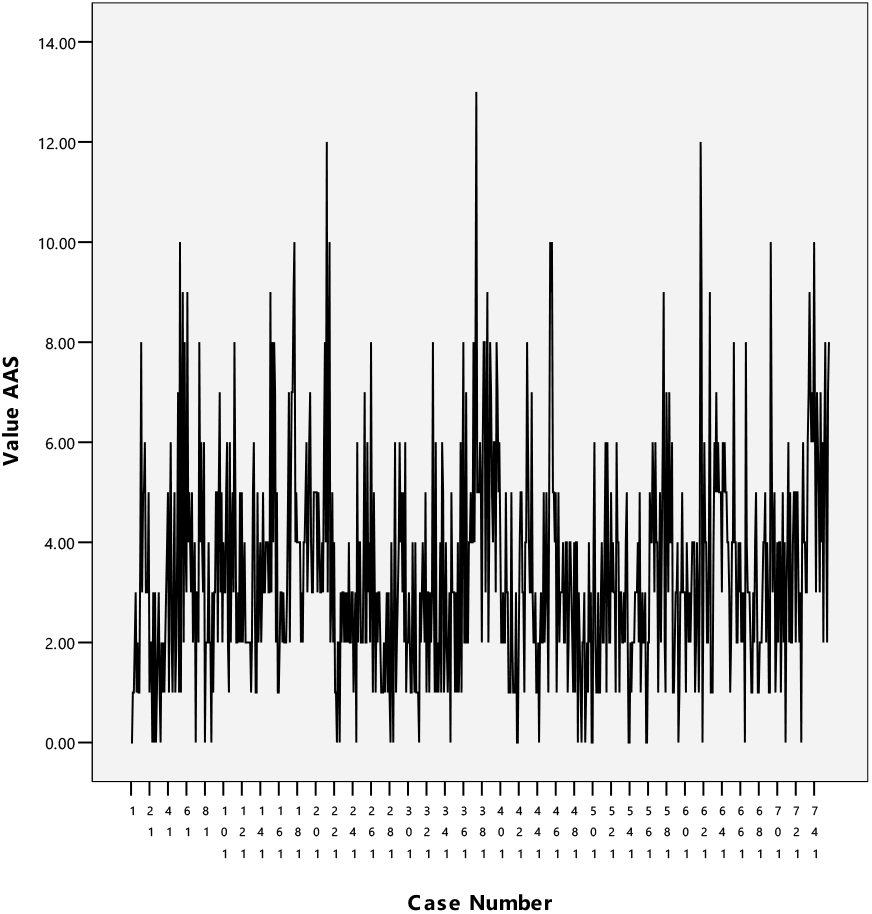
Site polymorphism of PB1.

**Table 2.** Spearman and pearson correlation of RV with RSA.

|  | Spearman ‘s rho | Pearson |
| --- | --- | --- |
| PB1 | 0.303 | 0.359 |
| NS5B a | 0.420 | 0.424 |

P<0.001 a ref (81)

**Fig. 3.**
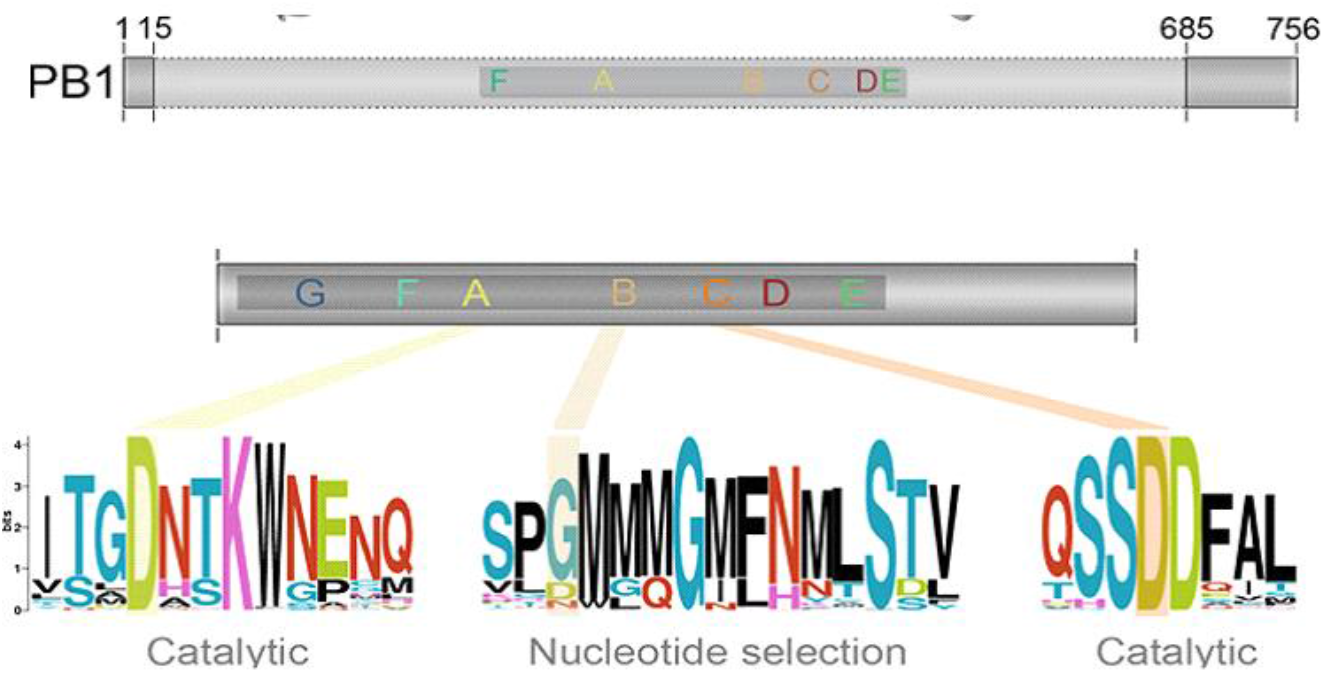
Schematic plot of PB1 varibility.

### ENTROPY

Average Shannon entropy values were also obtained for individual amino acid positions by using Bioedit Program. The resulting entropy values vary between 0 (total conservation) to 4.332 (complete variation). The mean value of less than 1.0 represent high conservation while less than 2.0 represent intermediate degree of amino acid conversation in the provided sequences(22). For IAV PB1, which gave a mean value of 0.077+_0.006, which is less than 1, indicates high conservation. Compares with HCV NS5B 0.422+_0.0216, which means more conservative than HCV.

Polymorphism and conservatism among site between group has a mean entropy value of 0.029+_0.004, showing that there is strong conservative of aa physical constraint.

**Fig. 4.**
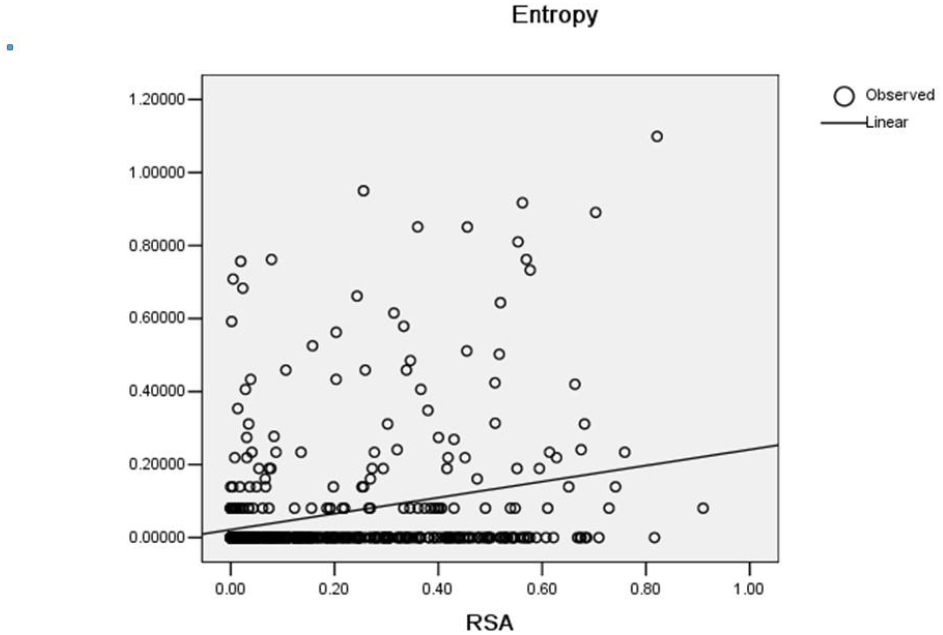
Liner correlation of aa entropy of PB1 with RSA.

**Fig.5.**
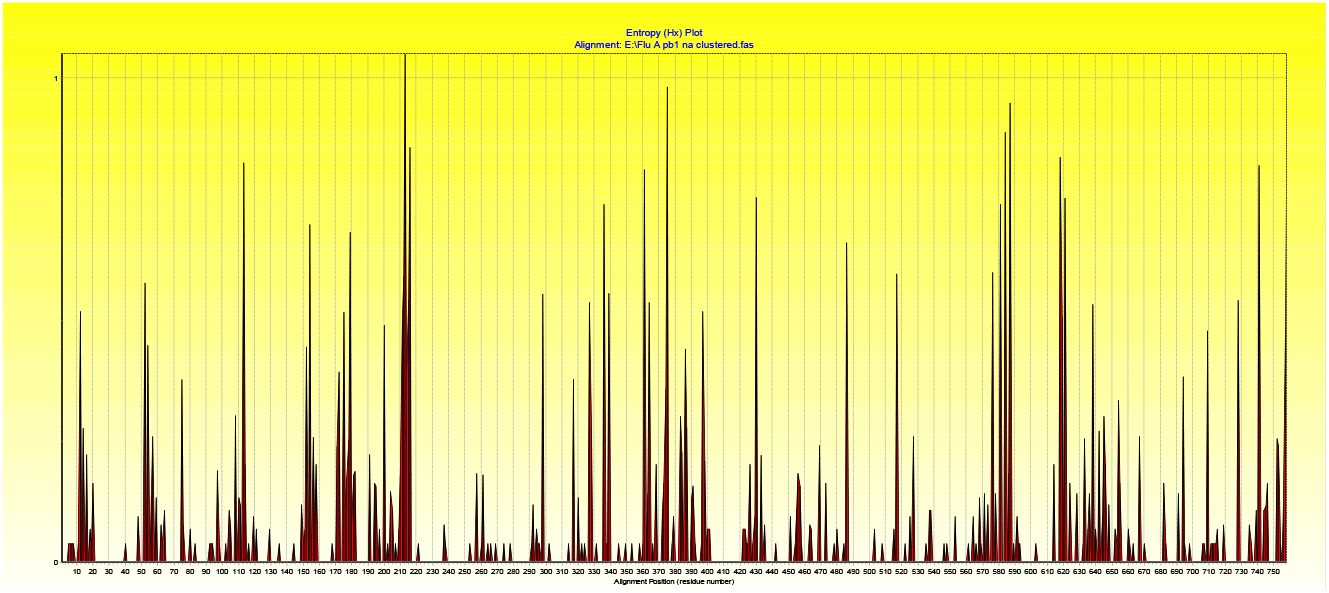
Site entropy of PB1.

### Ka/Ks

Classically purifying selection can be inferred by absence, for example, in the Ka/Ks test. We employ the normalized rate of occurrence of substitutions at synonymous sites (Ks) in a protein coding gene as a measure of the background rate of evolution, comparing this to the normalized rate of nonsynonymous changes. A dearth of the latter compared with the former (Ka/Ks < 1) is taken to imply that protein changing mutations happened but were too deleterious to persist (<u>Li et al. 1985</u>; <u>Goldman and Yang 1994</u>).

Our result gave HCV NS5B a mean value of Ka/Ks 0.095+_0.009, which is far less than 1 indicates strong purifying selection over all the protein sequences. Meanwhile, Ka/Ka increases with RSA, showing purifying selection decrease with aa expose to solvent. Compares with HCV NS5B 0.140+-0.012, which means IAV is more conservative than HCV. The relationship between solvent exposure and evolutionary rate (dN/dS) is found to be strong, positive, and linear. in accordance with Franzosa EA and Xia Y’s result and HCV HCV NS5B.

**Fig 6.**
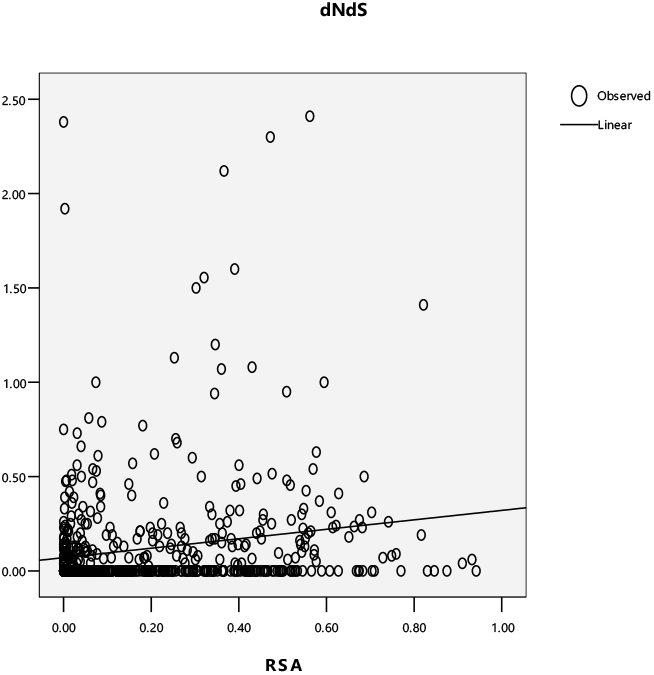
Liner correlation of dN/dS of PB1.

**Fig 7.**
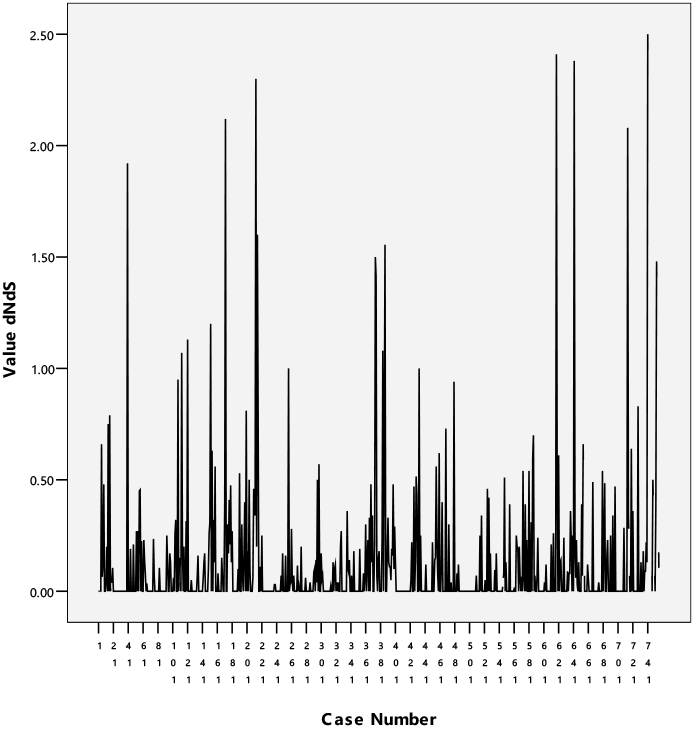
Site dN/dS of PB1.

### RNA secondary structure

**Fig.8.**
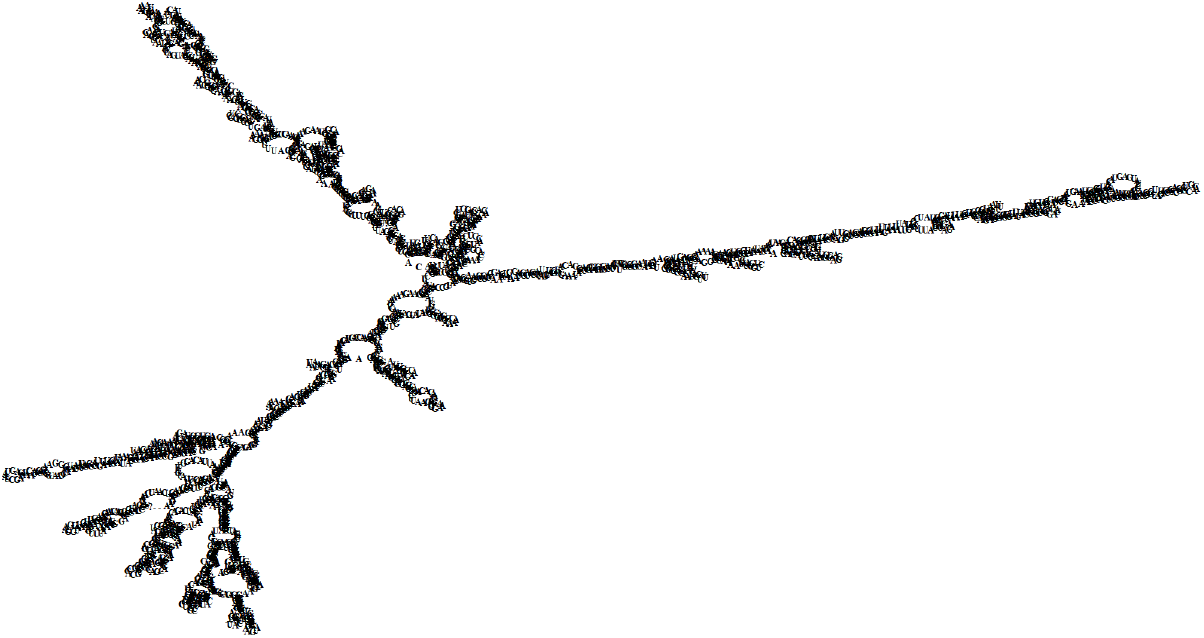
Schematic plot of secondary structure of PB1 coding RNA.

## Discussion

Sequence polymorphisms are fundamental features of all protein families, reflecting the dynamic interplay between mutation and selection. Our study of influenza A virus (IAV) PB1 protein reveals how structural constraints shape this evolutionary landscape. As an RNA virus with an error-prone polymerase (mutation rate: 1-8×10^−3^ substitutions/site/year), IAV exhibits greater genetic diversity than enteric bacteria but less than HCV, highlighting distinct evolutionary pressures across pathogens.

In this paper, we investigated the element that predict the relative likelihood that the site will be polymorphic within IAV PB1 protein. We found a strong positive relation between polymorphism and solvent accessibility, suggesting that amino acid sites that are more solvent-accessible are less likely be constrained in identity. This is the same with Bustamante CD’s result obtained from the study of the polymorphisms of enzymes in *E. coli* and *S. enteric* (1). This finding is in accordance with work done on multiple families of proteins showing that solvent accessibility impact amino acid substitution rates (25–29). This result is also in accordance with the works done on the proteins of yeast (30,31) and many virus proteins (32,33) especially HCV(81).

Of the 757 amino acids of this protein, only 33 aa are conservative, other 724 aa are polymorphic. Many functional residues like putative triad SDD (82) are polymorphic, as in the case of HCV NS5B for traid GDD. The 651P which is essencial for efficient RNA synthesis(82) are polymorphic. As suggested by Wellner A’s result, the laboratory enabled K219S exchange of PGK, which suggested given enough time and variability in selection levels, even utterly conserved and functionally essential residues might change (35). Further study needbe done to test this hypothesis.

Although many site-directed mutagenesis studies of several model proteins have demonstrated clearly the ability of protein secondary and 3D structures to accommodate extensive residue variability, the exact limits of the latter have remained a matter of conjecture. Kapp OH’s results (18) for the globin family provide reliable estimates of the maximum residue variability per position, 8-13 overall; furthermore, conservative estimates of residue variability at the interior positions vary tween 5 and 7, and those solvent-accessible positions appear to have very little if any restriction in substitution (residue variability 10-15). Our results show that for IAV PB1, this is also the case. Reliable estimates of the maximum residue variability per position, 13 overall, compare with NS5B of HCV, 18, which shows that IVA is less mutable than HCV(81). Mutation studies have showed that for most of sites of a protein, nearly all of the 20 amino acids can be replaced without significant influence of the function and structure of the protein (41–42,48–51). While in biological system, this is not the case, there is very strong constraints of residue variability at all sites, which show that many of the replacement may be slightly deleterious and are not be allowed.

Although there is considerable variability of the inner residues (residue maximum variability, 10), the inner part of IAV PB1 is very conservative with a mean of residue variability 3.0, (compare with globin family with 5 and HCV NS5B, 3.8). Furthermore, part of those solvent-accessible positions appear to have very little restriction in substitution, while others have strict constrains and remain monomorphic. Reliable estimates of the maximum residue variability position, 13 compares with HCV NS5B, 18 overall (81). These may all reflect that structural and functional constraints of this protein is very strong both in the inner part and at the surface and IAV is less variable than HCV.

### Apart from polymorphism, there are also conservatism of PB1 protein

Studies also showed that many surface strict conservative residues are involved in enzyme catalysis activity. There are 33 aa which are completely conservative, among them 17 aa are at the molecular surface (this 5% cut-off was devised and optimized by Miller et al. (34)), quite a lot of substitutions are the same physicochemical class, which means structural and functional constraints, shows that they are under strong selection pressure.

The PB1 protein contains the conserved motifs characteristic of RNA-dependent RNA polymerases and catalyzes the sequential addition of nucleotides during RNA chain elongation. The PB1 RNA polymerase domain harbours seven regions or motifs, which are arranged in the order G, F1–3, A, B, C, D and E from amino- to carboxy-terminus. The conserved motifs A and C in the palm subdomain contain the aspartate residues that bind to the two metal ions that coordinate nucleotide condensation. The active site is structurally conserved, whereas a clear divergence at the amino acid sequence level is present in motifs A and C of sNSV and nsNSV RNA polymerases. A priming loop protrudes into the active site cavity and is involved in the formation of the first dinucleotide during de novo initiation (37).

Of the 33 monomorphic sites, most of them are on the surface of PB1, only three sites are solvent inaccessible. Nearly all of the codons of these aa are polymorphic, suggesting that they are not constrained by virus RNA secondary structure formation (see discussion below). Apart from this, most of these sites are conservative in IBV, ICV, and IDV (data not shown). These all suggest that they are functional important. Further studies needed to test these possibilities.

This strict conservation of surface residue may also reflect that there are functional constraints of this protein by the interaction with other proteins. Various studies have showed that protein structure and sequences might be further constrained by selection for interactions among proteins (38). The trimeric viral RNA-dependent RNA polymerase, consisting of polymerase basic protein 1 (PB1), polymerase basic protein 2 (PB2) and polymerase acidic protein (PA) subunits, is responsible for the transcription and replication of the viral RNA genome segments. Biochemical studies indicated that the C terminus of PA interacts with the N terminus of PB1 and the C terminus of PB1 interacts with the N terminus of PB2, suggesting an N-terminal to C-terminal linear PA-PB1-PB2 arrangement of the subunits. However, three-dimensional images obtained by single particle analysis using electron microscopy indicate that the RNA polymerase forms a compact globular structure, suggesting more intimate interactions between all three subunits. High-resolution structural studies confirmed the interaction domains between the PA and PB1 as well as between the PB1 and PB2 subunits. As elucidated by the X-ray crystal structure, PA interacts extensively with PB1 with a total buried surface area of 17,330 Å. In contrast, PA only interacts marginally with PB2, and the total buried surface area between PA and PB2 is only 2,880 Å. The PB1 subunit interacts with the PB2 subunit extensively with a total buried surface area of 14,100 Å (37). For IAV PB1, many studies have showed that it has interaction with host proteins. Many of these strictly conservative surface residues may involve in these interactions.

Three key findings emerge about structural constraints:

1. Core conservation: Despite theoretical capacity for variation (max RV=10), buried residues show low variability (mean RV=3.0), stricter than globins (RV=5) or HCV NS5B (RV=3.8) (18,81).
2. Surface dichotomy: While some surface sites are hypervariable (RV≤13), others remain monomorphic, likely due to functional constraints from protein structure, function, folding and interactions (37).
3. Physicochemical filtering: 51% of conserved surface residues cluster in specific physicochemical classes (34), indicating purifying selection preserves functional properties.

Our entropy analyses confirm RSA’s dominant role:

- Shannon entropy strongly correlates with RSA (ρ=0.303, p<0.001)
- Grouped entropy decreases sharply with RSA, indicating physicochemical constraints
- Surface-core entropy differences mirror HCV NS5B (81), suggesting conserved evolutionary mechanisms

The Ka/Ks value of the whole protein indicate that there is a strong purifying selection in this protein, in accordance with previous studies (40). The correlation of Ka/Ks with RSA is used in conjunction with data on synonymous polymorphism to estimate quantitatively the reduction in purifying selection with increasing solvent accessibility. When compared with the distribution of synonymous polymorphism, the increased probability of amino acid polymorphism with solvent accessibility, suggesting strong purifying selection in areas of low solvent accessibility and weak purifying selection in areas of high solvent accessibility, irrespective of synonymy class, this result is the same with work down on *E*.*coli*. and *S. enteric* (1) and HCV NS5B(81).

The relatively high K_a_/K_s_ values of this protein reflect the high mutant rate of this protein and also the saturation property of its coding sequence at synonymous sites. For PB1, nearly all of the 3ed position of the coding sites are not saturated for both twofold and fourfold redundant sites(data not shown). There are saturation phenomena at synonymous sites in genes encoding protein families between species which may reflected strict constraints at those sites (58). For IAV PB1, the nonsaturation phenomenon may mean that purifying selection may be stronger than expected form the K_a_/K_a_ value. For Influenza virus, even nearly all of the 3ed position of the coding sites of IAV, IBV, ICV and IDV are not saturated (data not shown). Predicting the PB1 RNA secondary structure shown that nearly all of the PB1 coding RNA forms secondary structure which may constrains the muting of its RNA sequences and causes the nonsaturation of most of the synonymous sites. This may reflect that purifying selection at RNA sequences level is very strong.

Despite the new abundance of complete human influenza A virus genomes, little is known about adaptation outside the antigenic proteins mentioned above. Suzuki (60) applied a dN/dS approach to 100 complete H3N2 genomes and concluded that negative selection dominated in all proteins; significant positive selection was observed only in a handful of codons in the HA, NA, and nucleoprotein (NP) genes. Pond et al. analyzed the same data using a technique more sensitive to the detection of individual selective sweeps and reported more evidence for adaptation, in five genes: PB2 (polymerase basic 2-two codons), PB1 (polymerase basic 1-five codons), PA (polymerase acidic-three codons), HA (ten codons), and NA (four codons). Although the polymerase complex genes (PA, PB1, and PB2) often contained more adaptive substitutions than NP, M1, M2, and NS1, this appears to be an artifact of their greater length: per codon adaptation rates for the former are lower, indicating that positively selected sites in the polymerase genes are sparse

A subset of synonymous differences occurs at sites that are fourfold degenerate (that is, where a substitution with any base does not result in an amino acid replacement). As these sites are not subject to selective pressure at the protein level, base substitutions at many fourfold degenerate sites may accumulate rapidly. If influenza virus genes have been evolving in birds for long enough to reach evolutionary stasis, as is suggested by the high dS/dN ratios described above, one would predict that at many of the sites where fourfold degeneracy is possible, all four bases would be present in the avian clade unless the constraints of RNA secondary structure limit the accumulation of synonymous changes. In fact, when avian sequences from geographically distinct lineages (North American versus European) were compared, the percent difference at fourfold degenerate sites yielded values in the 27–38% range. In contrast, calculating the percent difference at fourfold degenerate sites in comparisons of the 1918 viral PA, PB1 and PB2 gene sequences with avian sequences yielded consistently higher values (range 41–51%) for all three genes. As with the other 1918 genes, this suggests that the donor source of the 1918 virus was in evolutionary isolation from those avian influenza viruses currently represented in the databases (79–80).

The K_a_/K_s_ value of two-fold redundant sites at the surface is smaller than that of proteins of *E. coli* and *S. enteric* (data not shown), this may reflect that purifying selection of PB1 may be stronger than that in *E. coli* and *S. enteric*, at least at the surface of PB1. This may be due to the reason that many of its surface residues are functional important and remain conservative; and there is strong purifying selection by the constraints of interaction of PB1 with other host proteins and other virus proteins. The trimeric viral RNA-dependent RNA polymerase, consisting of polymerase basic protein 1 (PB1), polymerase basic protein 2 (PB2) and polymerase acidic protein (PA) subunits, is responsible for the transcription and replication of the viral RNA genome segments. Biochemical studies indicated that the C terminus of PA interacts with the N terminus of PB1 and the C terminus of PB1 interacts with the N terminus of PB2, suggesting an N-terminal to C-terminal linear PA-PB1-PB2 arrangement of the subunits, without apparent direct interaction between PA and PB2. However, three-dimensional images obtained by single particle analysis using electron microscopy indicate that the RNA polymerase forms a compact globular structure, suggesting more intimate interactions between all three subunits. As elucidated by the X-ray crystal structure, PA interacts extensively with PB1 with a total buried surface area of 17,330 Å. The PB1 subunit interacts with the PB2 subunit extensively with a total buried surface area of 14,100 Å. These subunits interacting residues are submitted to severely constraint and remain conservative(37).

These constraints create an evolutionary paradox: while PB1’s error-prone polymerase generates diversity, structural and functional requirements maintain remarkable conservation. This balance reflects the polymerase’s dual needs - adaptability for host switching and stability for core functions. Our findings align with broader principles of protein evolution (73–78), demonstrating how structural biology can predict evolutionary trajectories across the tree of life.

